# Cell-type specific expression of oncogenic and tumor suppressive microRNAs in the human prostate and prostate cancer

**DOI:** 10.1101/251090

**Authors:** Binod Kumar, Avi Z. Rosenberg, Su Mi Choi, Karen Fox-Talbot, Angelo M. De Marzo, Larisa Nonn, W. Nathaniel Brennen, Luigi Marchionni, Marc K. Halushka, Shawn E. Lupold

**Author notes:** Financial Support: NIH (R01-CA143299, P30-CA006973, U42-OD011158), the Patrick C. Walsh Prostate Cancer Research Fund, the Department of Defense (W81XWH-10-2-0056, W81XWH-10-2-0046), the American Heart Association [13GRNT16420015], the Prostate Cancer Foundation Young Investigator Award, the Allegheny Health-Network-Hopkins Cancer Research Fund, and the Maryland Cigarette Restitution Fund. Corresponding Author: Shawn E. Lupold; 600 N. Wolfe St, Park 205, Baltimore, MD 21287-2101.

## Abstract

MiR-1 and miR-143 are frequently reduced in human prostate cancer (PCa), while miR-141 and miR-21 are frequently elevated. Consequently, these miRNAs have been studied as cell-autonomous tumor suppressors and oncogenes. However, the cell-type specificity of these miRNAs is not well defined in prostate tissue. Through two different microdissection techniques, and droplet digital RT-PCR, we quantified these miRNAs in the stroma and epithelium of radical prostatectomy specimens. In contrast to their purported roles as cell-autonomous tumor suppressors, we found miR-1 and miR-143 expression to be predominantly stromal. Conversely, miR-141 was predominantly epithelial. MiR-21 was detected in both stroma and epithelium. Strikingly, the levels of miR-1 and miR-143 were significantly reduced in tumor-associated stroma, but not tumor epithelium. Gene expression analyses in human cell lines, tissues, and prostate-derived stromal cultures support the cell-type selective expression of miR-1, miR-141, and miR-143. Analyses of the PCa Genome Atlas (TCGA-PRAD) showed a strong positive correlation between stromal markers and miR-1 and miR-143, and a strong negative correlation between stromal markers and miR-141. In these tumors, loss of miR-1 and gain of miR-21 was highly associated with biochemical recurrence. These data shed new light on stromal and epithelial miRNA expression in the PCa tumor microenvironment.

## INTRODUCTION

Comparative gene expression analyses between benign and malignant tissues have been critical to our current understanding of microRNAs (miRNAs) in human cancer (1). However, the vast majority of this data has been derived from macrodissected tissues, which contain both malignant and non-malignant cells. Consequently, miRNA expression from non-malignant cancer-associated cells may provide misleading clues for cancer cell biologists. For example, the tumor suppressive miR-143/145 cluster was recently found to be expressed and active in the stromal, rather than epithelial, compartment of the colon and lung (2,3). These new discoveries strongly suggest that the observed loss of miR-143/145 expression in colon and lung carcinomas has been due to differential stromal sampling between benign and malignant tissues, rather than from the loss of miRNA expression within cancer cells. These results challenge the role of miR-143/145 as a cell-autonomous tumor suppressor, and instead, indicate a functional role within the tumor microenvironment (4). In light of this, there is an urgent need to characterize the cell-type specific expression of many miRNAs in normal human tissues, within cancer cells, and within the tumor microenvironment.

MiRNA expression can be directly analyzed in specific cell types, or tissue compartments, by *in situ* hybridization. However, transcript size, stability, and expression level have made this challenging in most clinical specimens. Laser capture microdissection (LCM) offers an alternative method to select and capture specific cell types for subsequent RNA extraction and miRNA quantification (5). Expression microdissection (xMD) is another developing cellular/subcellular isolation method that provides high throughput, and operator-independent, selection and capture of cells for downstream analyses (6). Similar to LCM, xMD applies illumination-based membrane melting and cell capture, but through a semi-automated approach where immunohistochemical (IHC) staining with dark chromogens and whole-slide irradiation are applied to locally heat and capture all strongly-stained cells from a single slide or section. Here we apply both xMD and LCM to evaluate miRNA gene expression patterns in the stromal and epithelial compartments of normal and malignant prostate tissue.

Prostate cancer (PCa) is one of the leading causes of cancer death in American men (7). Gene expression analyses from macrodissected radical prostatectomy specimens have uncovered several potential oncogenic and tumor suppressive miRNAs (8). Four miRNAs have been frequently reported to have aberrant expression in PCa: miR-1, miR-21, miR-141, and miR-143. The levels of miR-1 and miR-143 are commonly diminished in PCa, and ectopic expression of either miR-1 or miR-143 inhibits the growth and survival of PCa cells (9–13). Thus, miR-1 and miR-143 have been considered cell-autonomous tumor suppressors of PCa. Conversely, miR-21 and miR-141 levels are frequently elevated in human PCa (14–17). Ectopic expression of miR-21 enhances PCa cell proliferation and tumor growth, and it imparts therapeutic resistance, while ectopic over-expression of miR-141 enhances PCa cell survival and suppresses stemness (16,18–21). Thus, miR-21 and miR-141 have been considered as cell-autonomous drivers of PCa cell growth and survival. Here, to further investigate these four miRNAs, we applied xMD and LCM to analyze the stromal and epithelial expression of each miRNA in benign and malignant prostate tissue. Our results reveal predominant miRNA expression patterns in specific cell types, and they uncover novel miRNA expression changes in the tumor microenvironment, which were previously believed to occur within PCa cells.

## MATERIALS AND METHODS

### Immunohistochemistry

IHC was performed using standard methods. Radical prostatectomy specimens were obtained from the Johns Hopkins Hospital Pathology Department. Sample use required informed consent and internal review board permission. Slides were reviewed by a genitourinary pathologist (ADM) and four cases that included a large localized region of tumor and a separate section of uninvolved prostate were selected. Slides were cut from FFPE blocks at 4 microns, using RNA-precautions, and kept desiccated in Drierite. Slides were baked 20 minutes at 60 °C, paraffin was removed by immersion in 2 changes of xylene, graded ethanol washes, and rehydration in water. Antigen retrieval was performed by immersion in Trilogy (Sigma-Aldrich) at 126 °C/18-23 psi. Slides were stained for AE1/AE3 (Diagnostic Biosystems) or α-SMA (Abcam) followed by polymer HRP IgG (Leica Biosystems). Protector RNase Inhibitor (Roche Diagnostics) was added to antibody incubations (1 U/µL). The antibody complex was detected with Deep Space Black (Biocare Medical).

### Microdissection

xMD was performed as previously described (22). Briefly, the region of interest was overlaid with ethylene vinyl acetate (EVA) polymer film (3M) and vacuum sealed (FoodSaver) to approximate film and tissue. Slides were light-flashed 5X at power 5 by a SensEpil flash lamp (Silk’n) (Supplementary Figure S1). The membrane was removed, microscopically dissected to remove adjacent inappropriate tissue, and stored at ࢤ80 °C. Digital images were taken using an AMA40 camera adapter (AmScope), digital camera, and BH2 light microscope (Olympus). For xMD, seven consecutive sections were isolated for each sample and combined for RNA extraction. For LCM, prostate stroma and epithelium from benign areas was collected as described earlier by Nonn and colleagues (23,24). Briefly, formalin-fixed paraffin-embedded specimens were deparaffinized, fixed, stained (0.5 % toluidine blue) and microdissected using a LMD-6000 LCM instrument (Leica). Stroma and epithelium from surrounding benign regions were collected into Eppendorf caps containing digestion buffer (Thermo Fisher).

### miRNA isolation and quantification

Total RNA from normal and primary PCa tissue was obtained from Colm Morrissey (University of Washington, Seattle) and the PCBN. RNA was isolated from scraped tissue or EVA membranes by Recover All Total Nucleic Acid Isolation Kit (Life Technologies). Cel-miR-39 was spiked into each xMD sample after digestion. RNA quantification was performed by Qubit HS RNA kit (Life Technologies). TaqMan MicroRNA RT (Applied Biosystems, Life Technologies) was used to generate cDNA for hsa-miR-21-5p, hsa-miR-143-3p, hsa-miR-141-5p, hsa-miR-1-5p, cel-miR-39, and RNU6B from 10 ng of RNA. Droplet Digital PCR (ddPCR) (Bio-Rad QX200) was performed using 1.5 µl of cDNA. A minimum of 10,000 droplets was considered as a positive read out. All results were normalized per 10,000 copies of cel-39 for xMD samples, or 10,000 copies of RNU6B for non-xMD samples.

### Cell line and stromal cell culture

Six primary stromal cultures were derived from healthy and cancerous prostate tissue in accordance with IRB-approved protocols. Healthy prostate was defined as tissue from young men (< 25 yo) obtained through a rapid organ donor program organized by the National Disease Research Interchange. Tissue was perfused, surgically harvested, and delivered within 24 hrs of the time of death. PCa tissue was obtained from radical prostatectomy patients treated at the Brady Urological Institute at Johns Hopkins via the Prostate Biospecimen Repository. Tissue was dissociated into a single cell suspension and cultured in Rooster High Performance Media (RoosterBio) with regular media changes every 3-4 days as previously described (25,26). LNCaP PCa cells (ATCC), PrEC normal prostate epithelial cells (a gift from William Isaacs, Johns Hopkins), BPH1 cells (a gift from John Isaacs, Johns Hopkins), HFF Human Foreskin Fibroblasts (William Isaacs), and human fibroblasts derived from coronary artery cells were maintained in RPMI 1640 supplemented with 10 % Fetal Calf Serum. All cells were maintained in a 5 % CO_2_, 95 % air humidified incubator at 37 ˚C and split at ≤ 80 % confluency.

### Western blot analyses

Equal amount of proteins (20 μg) were separated by SDS-PAGE, transferred to nitrocellulose membrane, and immunoblotted using E-cadherin (3195) antibodies from Cell signaling, or Vimentin (sc-32322) or CK18 (sc-6259) antibodies from Santa Cruz Biotechnology, or α-SMA (ab7817) from Abcam, or β-actin (A5441) or GAPDH (G9545) from Sigma. Bands were detected and quantified with IRdye 800CW (926-322210) or IRdye 680LT (926-68021) by LI-COR odyssey imaging system and software (LI-COR Biosciences).

### Statistical and bioinformatic analyses

Comparative analyses of clinical samples applied Mann-Whitney Wilcox rank-sum and cell culture response analyses applied student’s t-test. Results were analyzed by GraphPad Prism (GraphPad Software) and significance was assigned with p < 0.05. Normalized microRNA expression data were retrieved from the UCSC Xena Functional Genomic Browser and analyzed using R-Bioconductor packages. Survival analysis was performed by univariate and multivariate Cox proportional hazards model, using time to biochemical recurrence as the outcome. MicroRNA expressions were used as continuous and dichotomized values, using optimal thresholds identified by maximizing the log-rank test (27). Pearson’s correlation analysis was performed by linear regression with lines fitted using the least-squares method.

## RESULTS

### miR-21 and miR-141 levels are elevated in human PCa, and miR-1 and miR-143 levels are reduced

The expression of miR-21 (hsa-miR-21-5p) and miR-141 (hsa-miR-141-5p) are frequently reported to be elevated in human PCa, while the levels of miR-1 (hsa-miR-1-5p) and miR-143 (hsa-miR-143-3p) are frequently found to be decreased (8). To confirm these observations, we quantified the levels of each miRNA in normal and malignant human prostate tissue by reverse transcription and droplet digital PCR (RT-ddPCR). This approach has been demonstrated to provide precise miRNA quantification across a broad array of concentrations (28). Total RNA from macrodissected normal prostate (n = 15) and primary PCa (n = 15) were obtained from the University of Washington through the Prostate Cancer Biorepository Network (PCBN). Mature miRNA quantification, normalized to the stable internal control RNU6B (29), verified the expected expression patterns for each gene (Figure 1A). Specifically, miR-1 and miR-143 levels were significantly lower in PCa, by 2.3 fold (p = 0.025) and 1.4 fold (p = 0.025), respectively, when compared to normal prostate. Conversely, miR-141 and miR-21 levels were significantly higher in PCa by 1.5 fold (p = 0.01) and 1.9 fold (p = 0.025). The normalized copy numbers are presented on a log scale for comparison of all four miRNAs on a single graph.

**Figure 1.**
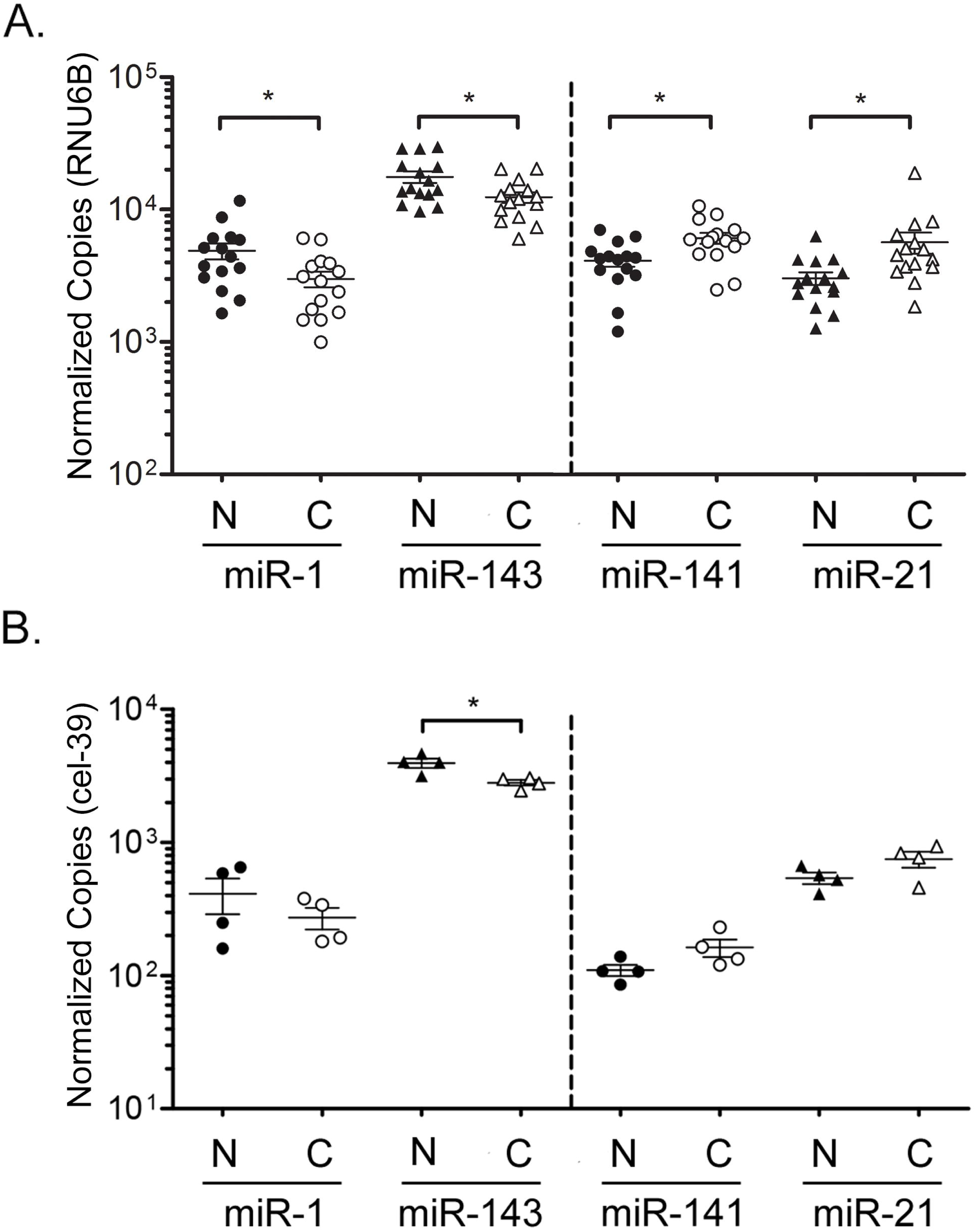
Aberrantly expressed miRNAs in human prostate cancer. Dashed line separates miRNAs decreased in cancer versus increase in cancer. **A**. Differential levels of miR-1, miR-143, miR-141, and miR-21 in macrodissected normal (N) and malignant (C) prostate tissue from the University of Washington (n = 15 per group) by RT-ddPCR. Copy numbers normalized per 10,000 RNU6B. **B**. RT-ddPCR quantification of miRNAs in a separate set of macrodissected normal and malignant prostate tissue from Johns Hopkins radical prostatectomy (n = 4 per group). Copy numbers normalized per 10,000 copies cel-39 internal control. These sample are applied to xMD in Figures 2 and 4. Bars represent mean and standard error. Log Scale. P-values determined by Wilcox Rank Sum analysis. *, p < 0.05.

Four separate cases of low-to-intermediate risk primary PCa (Supplementary Table S1) were then selected from Johns Hopkins University (JHU) radical prostatectomy tissue blocks for microdissection. Prior to microdissection, pathologist-defined whole benign and malignant tissue from each case were macrodissected and total RNA was isolated. A non-human miRNA, cel-miR-39 (cel-39), was spiked into each tissue sample during extraction as an internal reference control for normalization. Similar trends of elevated miR-21 and miR-141, and reduced miR-1 and miR-143, were found in these cases (Figure 1B).

### Compartmentalized expression of miR-1, miR-141, and miR-143 in normal human prostate

Adjacent non-malignant regions from the same JHU radical prostatectomy cases (Figure 1B) were then microdissected by xMD, to isolate the stromal and epithelial cells from normal prostate tissue (Figure 2A and Supplementary Figure S1). Specifically, stromal cells were isolated by anti-α-smooth muscle actin (α-SMA) IHC staining, with Deep Space Black Chromogen, followed by laser illumination and membrane-based cell capture (Figure 2B). Epithelial cells were similarly isolated by pan-cytokeratin IHC staining (AE1/AE3), and xMD cell capture. Mature miRNA copy numbers were then quantified by RT-ddPCR, and normalized to cel-39 spiked control. Strikingly, the expression of miR-1 and miR-143 were found to be almost exclusively stromal, with nearly undetectable levels in the epithelium. An average of 395 copies of miR-1 were detected in prostate stroma, while less than 10 copies could be detected in epithelium (Figure 2C). MiR-143 was more highly expressed in stromal cells, with an average of 526 copies, while only 10 copies were detected in the epithelium. In summary, miR-1 and miR-143 levels were 48.7 fold (p = 0.029) and 50.4 fold (p = 0.029) higher in the stroma, when compared to epithelium. Conversely, miR-141 was almost exclusively detected in epithelium, with an average of 45 epithelial copies and only 5 stromal copies. Thus, miR-141 was 7.8 fold (p = 0.029) higher in the epithelium, when compared to the stroma. MiR-21 was detected in both stroma and epithelium, with an average of 448 stromal copies and 25 epithelial copies. Levels of miR-21 were unexpectedly higher in the stroma, at 18.2 fold (p = 0.029).

**Figure 2.**
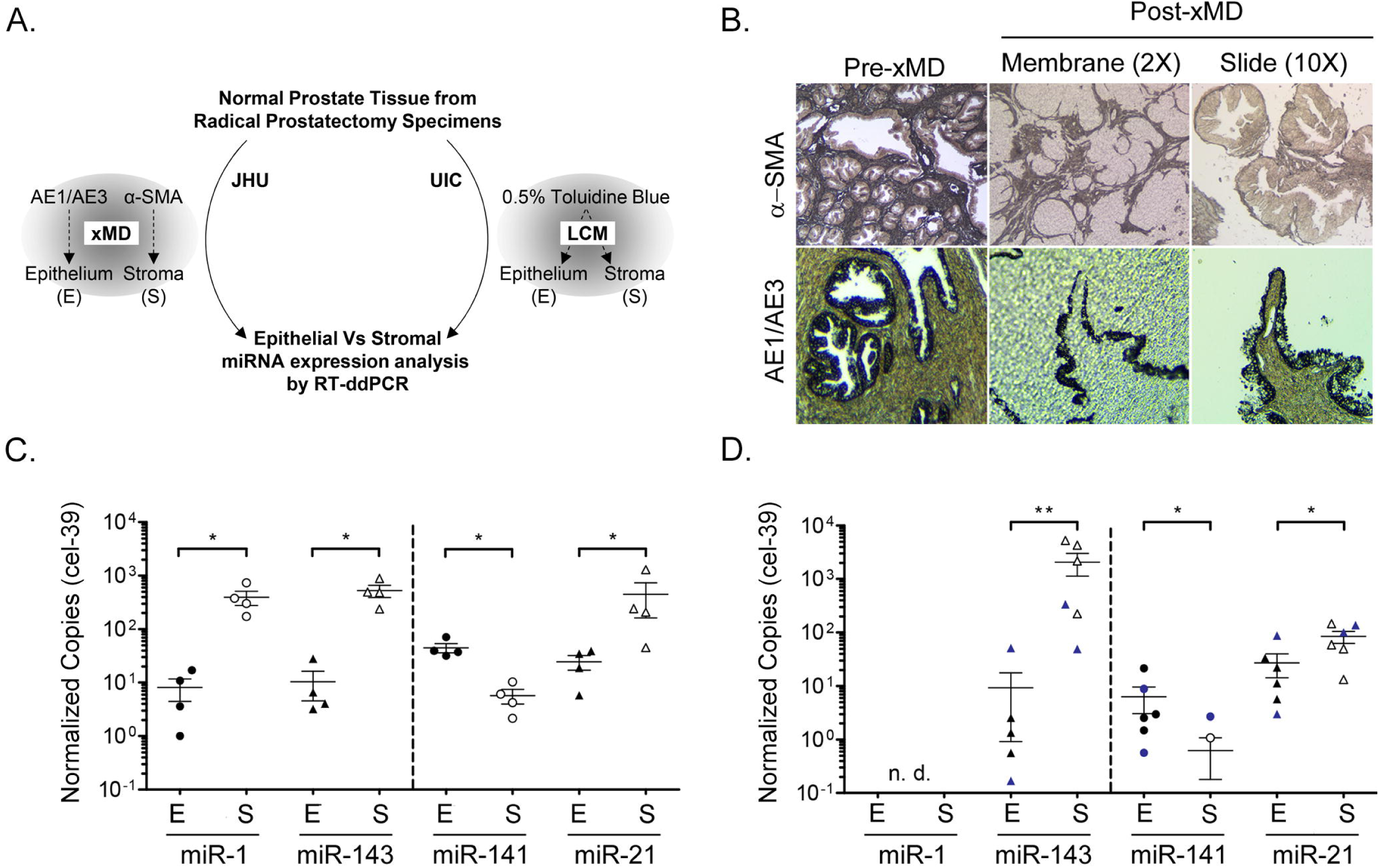
Compartmentalized expression of miR-1, miR-141, and miR-143 in normal human prostate tissue. **A**. Schematic of microdissection approaches applied to isolate prostatic stroma and epithelium from two separate sample sets. Johns Hopkins University (JHU). University of Illinois at Chicago (UIC). **B**. Representative images of slides and membrane before and after α- SMA and AE1/AE3 xMD. Magnification, 2× or 10×. **C**. RT-ddPCR quantification of miRNAs in xMD dissected prostate stroma (S) and epithelium (E) from JHU samples applied to Figure 1B (n = 4). Dashed line separates miRNAs decreased in cancer versus increase in cancer. **D**. RT-ddPCR quantification of miRNAs in LCM dissected prostate stroma (S) and epithelium (E) from normal prostate tissues microdissected at the UIC (n = 6). Blue symbols indicate patients who received Vitamin D. Bars represent mean and standard error; n.d., not detected. Log Scale. P-values determined by Wilcox Rank Sum analysis. *, p < 0.05; **p < 0.01.

To further investigate these results, miRNA levels were quantified in a separate set of normal prostate tissues, which were microdissected by LCM at the University of Illinois (Figure 2A, LCM). Total RNA had been previously isolated from normal prostate stroma and epithelium (n = 6) as part of a clinical trial involving oral Vitamin D3 (24). These specimens did not include an internal control, therefore miRNA copies are normalized to RNU6B. Surprisingly, miR-1 could not be detected in the stroma or epithelium of these samples (Figure 2D). Mir-143 was verified to be predominantly stromal in these tissues, with 222 fold (p = 0.004) greater expression in the stroma when compared to the epithelium. More specifically, 2,067 copies of miR-143 were detected in the stroma, while only 9 copies were detected in the epithelium. The predominant epithelial expression of miR-141 was also verified in these samples (10 fold, p = 0.026), with an average of 6.3 copies detected in epithelium and only 0.6 copies detected in stroma. MiR-21 was again detected in both tissue compartments with an average of 84 stromal copies and 27 epithelial copies. Again, levels of miR-21 were significantly higher in the stroma of these samples, but with a much lower fold difference (3.1 fold, p = 0.041). Two of these tissue samples were derived from patients who received oral Vitamin D3 (Figure 2D, indicated in solid blue). No general relationship between Vitamin D3 treatment and miRNA expression levels could be determined from these two samples, although previous analyses in larger sample sets have found that miR-21 levels are inversely associated with serum Vitamin D3 levels (24).

### Expression of miR-1, miR-21, miR-141, and miR-143 in human cells and tissues

To gain a better understanding of the cell-type specificity of these miRNAs, we examined miRNA expression levels across a series of cells and tissues from the RNA sequencing data published within the public Sequence Read Archive (SRA) (30). Cell and tissue data were grouped according to tissue origin and relative reads per million (RPM) were applied for comparisons. For each miRNA, the tissues were ranked from lower to higher expression levels. The levels of miR-1 were low (average < 10^3^ RPM) in most tissues, with the highest levels found in skeletal muscle and heart (Figure 3A). Distinctly, miR-1 expression was extremely low or absent (0 - 1 RPM) from fat and fibroblast samples. The levels of miR-141 were also low (< 10^3^ RPM) in most tissues. MiR-141 expression was lowest in muscle and heart tissues, and highest in epithelial-rich tissues such as skin, thyroid, and pancreas (Figure 3B). MiR-143 was detected at high levels (10^4^ – 10^6^ RPM) in most tissues (Figure 3C). Only one group, immune cells, expressed average levels below 10^3^ RPM. Like miR-143, miR-21 expression was also high in most tissues (10^4^ – 10^6^ RPM). No tissues displayed an average level lower than 10^3^ RPM. Immune, adrenal, lung, and fibroblast displayed the highest miR-21 expression levels (Figure 3D). These results show a generally tissue-restricted expression pattern for miR-1 and miR-141, and a broader and higher level expression pattern for miR-143 and miR-21, across multiple tissue types.

**Figure 3.**
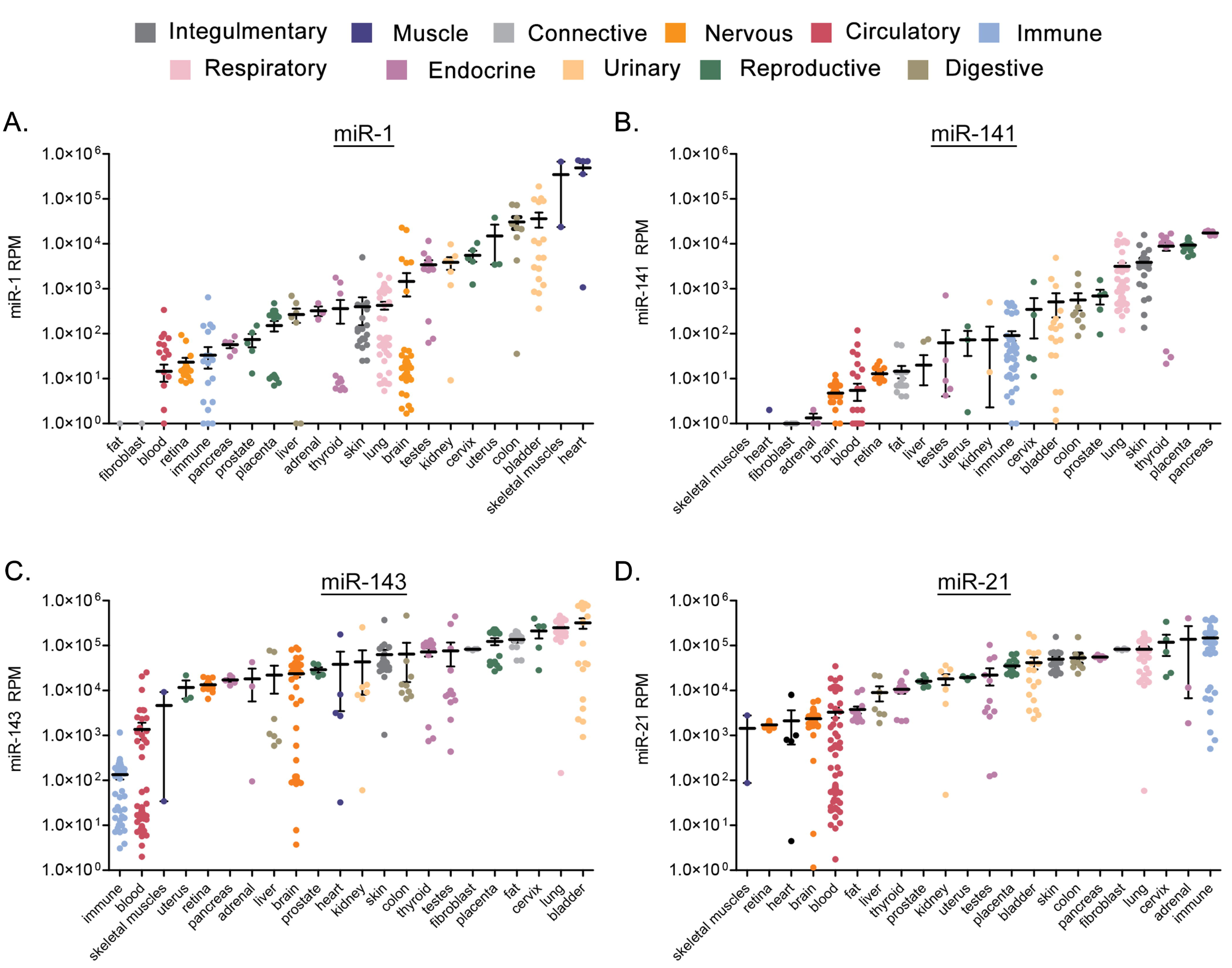
Cell and tissue type specific expression of miR-1, miR-143, miR-141, and miR-21. Expression levels from RNA sequencing data within the public Sequence Read Archive (SRA). Cell and tissue data were grouped according to tissue origin (Legend). Tissues ranked left to right from lower to higher expression levels per miRNA. **A**. miR-1, **B**. miR-141, **C**. miR-143, **D**. miR-21. Bars represent mean and standard error. Log scale.

### Levels of miR-1 and miR-143 are significantly reduced in tumor-associated stroma

Predominant stromal or epithelial miRNA expression could confound the interpretation of miRNA expression or function in adenocarcinomas, such as PCa, which exhibit higher epithelial cell densities, and lower stromal cell densities, when compared to adjacent normal tissue. To provide more focus on the cell-type specificity of miRNA expression in human prostate tumors, we microdissected the stromal and epithelial compartments from the cancerous regions of the same four JHU radical prostatectomy specimens applied to Figure 2 (Figure 4A, schematic). We then compared the levels of each miRNA in normal versus cancer stroma, and normal versus cancer epithelium. Notably, the levels of miR-1 were significantly lower (9.5 fold, p = 0.029) in tumor-associated stroma, when compared to normal prostate stroma (Figure 4B, left). However, the levels of miR-1 were not significantly different between normal and cancerous epithelium. The levels of miR-143 were also significantly lower in tumor associated stroma, by 5.4 fold (p = 0.029), when compared to normal prostate stroma (Figure 4B, right), while the levels of miR-143 were not significantly different between normal and cancerous epithelium. Significant differences could not be detected between normal and cancerous epithelium, or stroma, for miR-141 or miR-21 (Figure 4C). Therefore, similar conclusions of stromal or epithelial expression changes could not be made for these miRNAs.

**Figure 4.**
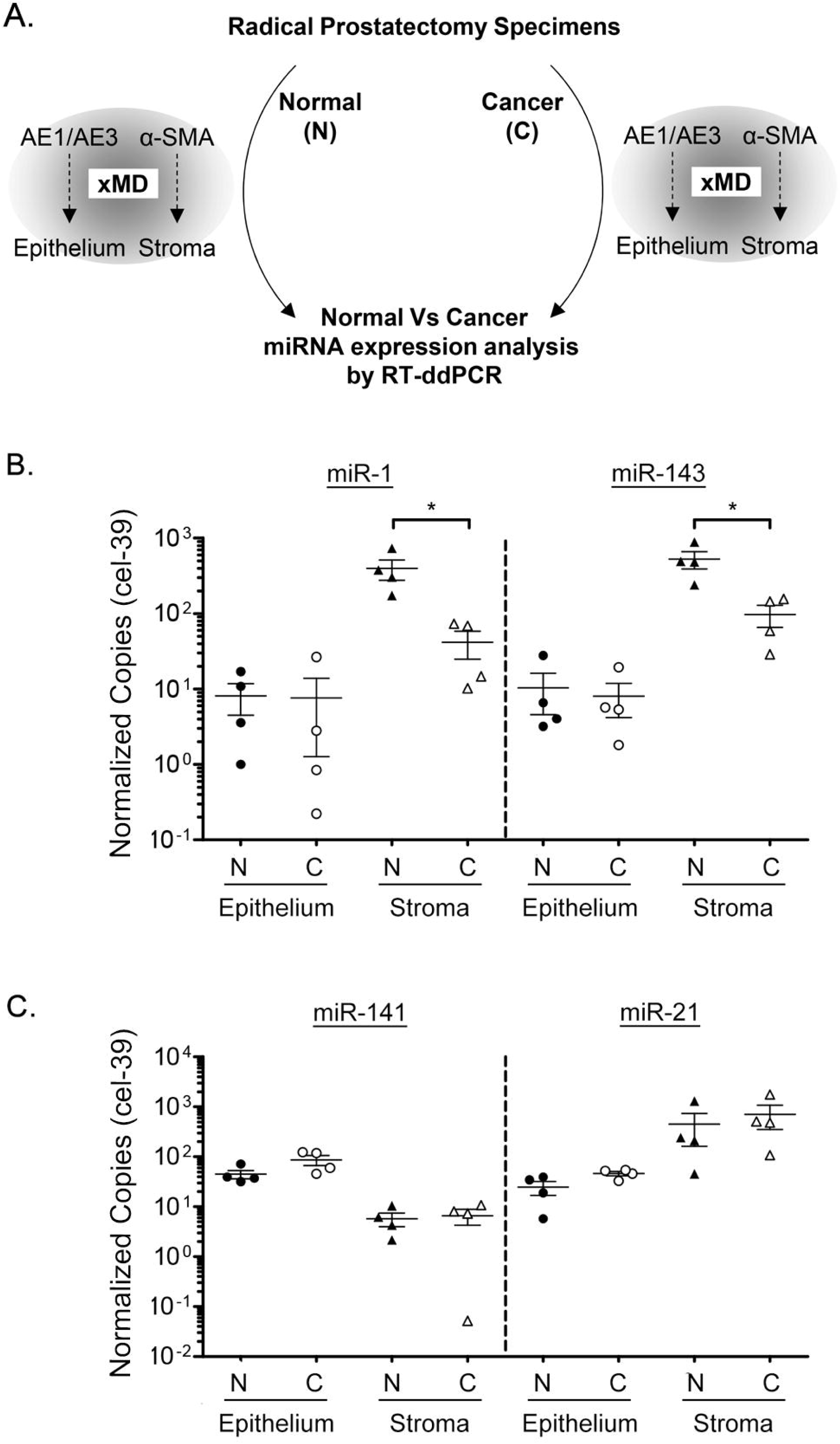
Reduced miR-1 and miR-143 levels in PCa-associated tumor stroma. **A**. Schematic of normal (N) versus cancer (C) comparison between miRNA levels in xMD microdissected stroma versus epithelium. Stroma was isolated by α-SMA xMD, and epithelium isolated by AE1/AE3 xMD, using JHU samples from Figure 1B (n = 4). Levels of each miRNA quantified by RT-ddPCR and copy number normalized per 10,000 cel-39. **B**. miR-1 and miR-143 levels in normal versus malignant epithelium and stroma. **C**. miR-141 and miR-21 levels in normal versus malignant epithelium and stroma. Bars represent mean and standard error. Log Scale. P-values determined by Wilcox Rank Sum analysis. *, p < 0.05

### Cell-type Specific miRNA expression in human prostate cell lines and stromal cell cultures

To further investigate the expression of these miRNAs in pure prostate epithelial cell populations, the level of each miRNA was quantified in normal prostate epithelial cells (PrEC), benign prostatic hyperplasia epithelial cells (BPH1), and LNCaP prostate adenocarcinoma cells (Figure 5A). Normal fibroblast cells, derived from primary coronary artery, were included as a reference control for stromal gene expression. Remarkably, we were not able to detect miR-1 in any of these cell extracts. MiR-143 was only detected at appreciable levels in fibroblasts, with extremely low levels detected in PrEC (0.03 copies) and BPH1 cells (0.005 copies). MiR-143 was undetectable in LNCaP cells. MiR-141 was detected at high levels in LNCaP, BPH1, and PrEC cells, while levels in fibroblast cells were extremely low (< 0.001 copies). Finally, miR-21 was detected at appreciable levels in all four cell lines.

**Figure 5.**
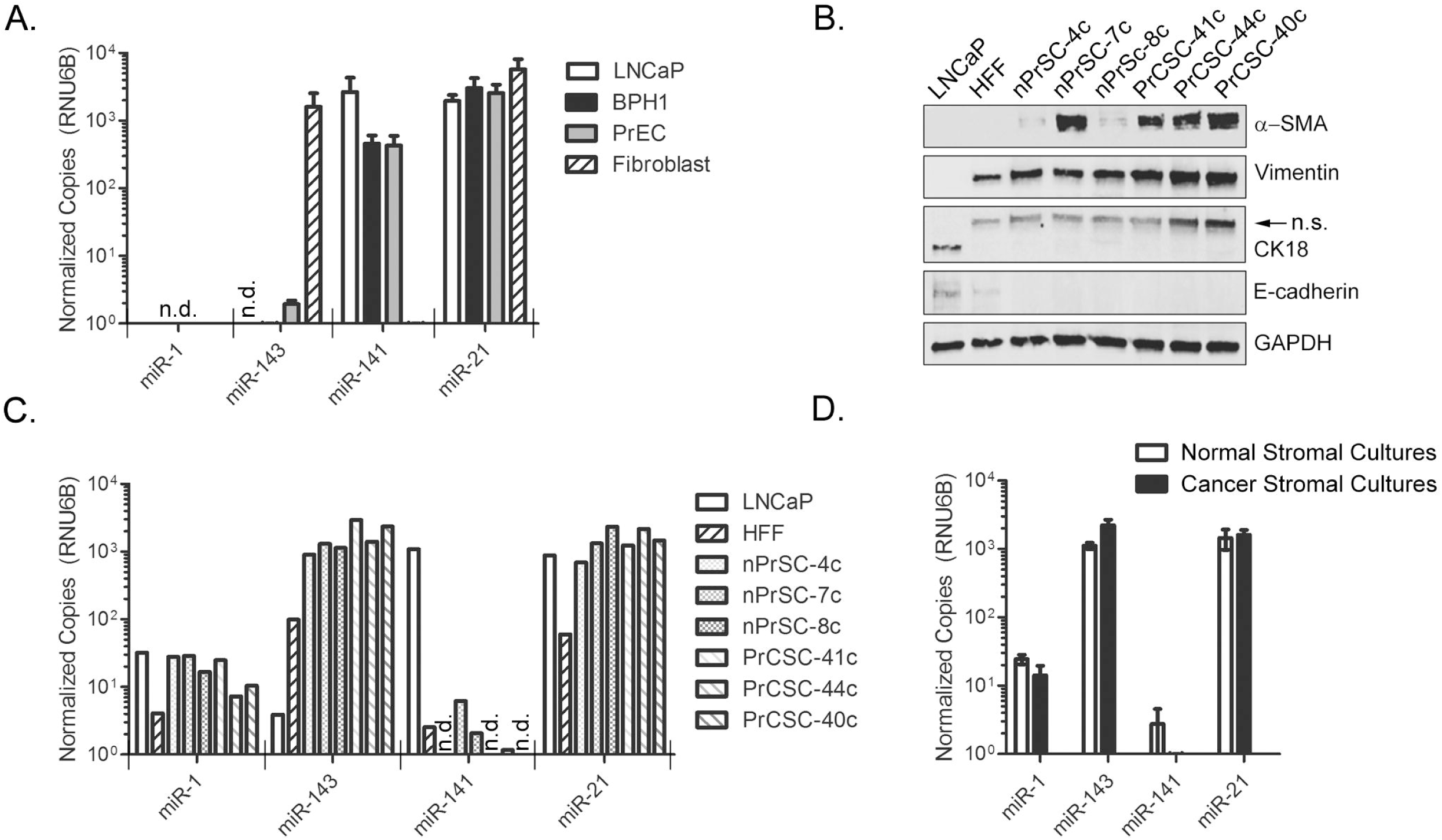
Cell-selective miRNA expression in prostate cell lines and stromal cultures **A**. RT-ddPCR quantification of miRNAs in epithelial prostate, BPH, and PCa cells lines and normal fibroblast cells. Results are mean normalized copies per 10,000 RNU6B, plus standard error from two independent experiments. miR-1 not detected. **B**. Characterization of stromal cultures derived from normal prostate (nPrSC-4c, nPrSC-7c, and nPrSC-8c) or from PCa (PrCSC-41c, PrCSC-44c, and PrCSC-40c) by WB. LNCaP PCa cells and Human Foreskin Fibroblasts (HFF) were included as reference controls for epithelial and stroma cells, respectively; n.s, non-specific. **C**. RT-ddPCR quantification of miRNAs in stromal cultures derived from normal prostate or from PCa. **D**. Comparison of miRNA copy number from normal stromal cultures (n = 3) versus PCa stromal cultures (n = 3). Bars represent mean plus standard error. No significant differences detected, Wilcox Rank Sum analysis. MiRNA copy numbers normalized per 10,000 copies RNU6B. Log scale.

We then studied the expression of these miRNAs in prostate stromal cultures, derived from three different normal human prostates (nPrSC-4c, nPrSC-7c, and nPrSC-8c), and from three different PCa tissue specimens (PrCSC-41c, PrCSC-44c, and PrCSC-40c). Each primary culture was characterized by western blotting for stromal and epithelial specific markers. Human PCa cells (LNCaP) and human foreskin fibroblast cells (HFF) were included as reference controls. All stromal-derived cultures were positive for vimentin and α-SMA, and negative for epithelial markers CK18 and E-cadherin (Figure 5B). MiR-1 was detectable at low copy number (7 - 29 copies) in these six stromal cultures (Figure 5C). Low miR-1 copy numbers could also be detected in these LNCaP (32 copies) and HFF (4 copies) cell extracts. MiR-143 was detected at high levels (905 – 2,930 copies) in all stromal cultures, as well as HFF cells (99 copies). Extremely low copy numbers of miR-143 were also detected in these LNCaP cell extracts (3.9 copies). The levels of miR-141 were extremely low (0 - 6 copies) in stromal cultures, when compared to LNCaP cells (1,092 copies). MiR-21 was detected in all cells, ranging from 693 to 2,993 copies in prostate stromal cultures. Lower levels of miR-21 expression (58 copies) were detected in HFF cells. Finally, we compared the mean expression of each miRNA in normal versus cancer derived stromal cultures (Figure 5D). No significant differences in miRNA expression were found between these *ex vivo* cultures.

### Cell-type specific miRNA expression in the PCa Genome Atlas (TCGA-PRAD)

The PCa Genome Atlas (TCGA-PRAD) is a valuable and publically available dataset that provides both mRNA and miRNA gene expression data from a large number of well-characterized patient samples. This dataset was derived from macrodissected tissue specimens, and therefore includes information from the epithelial and stromal compartments. Consistent with our analyses in human tissue, we found increased levels of miR-21 (1.1 fold, p < 0.001) and miR-141 (1.3 fold, p < 0.001) in PCa cases, and decreased levels of miR-143 (2.6 fold, p < 0.001) (Figure 6A). The levels of miR-1 were not significantly different between benign and malignant tissue in this dataset.

**Figure 6.**
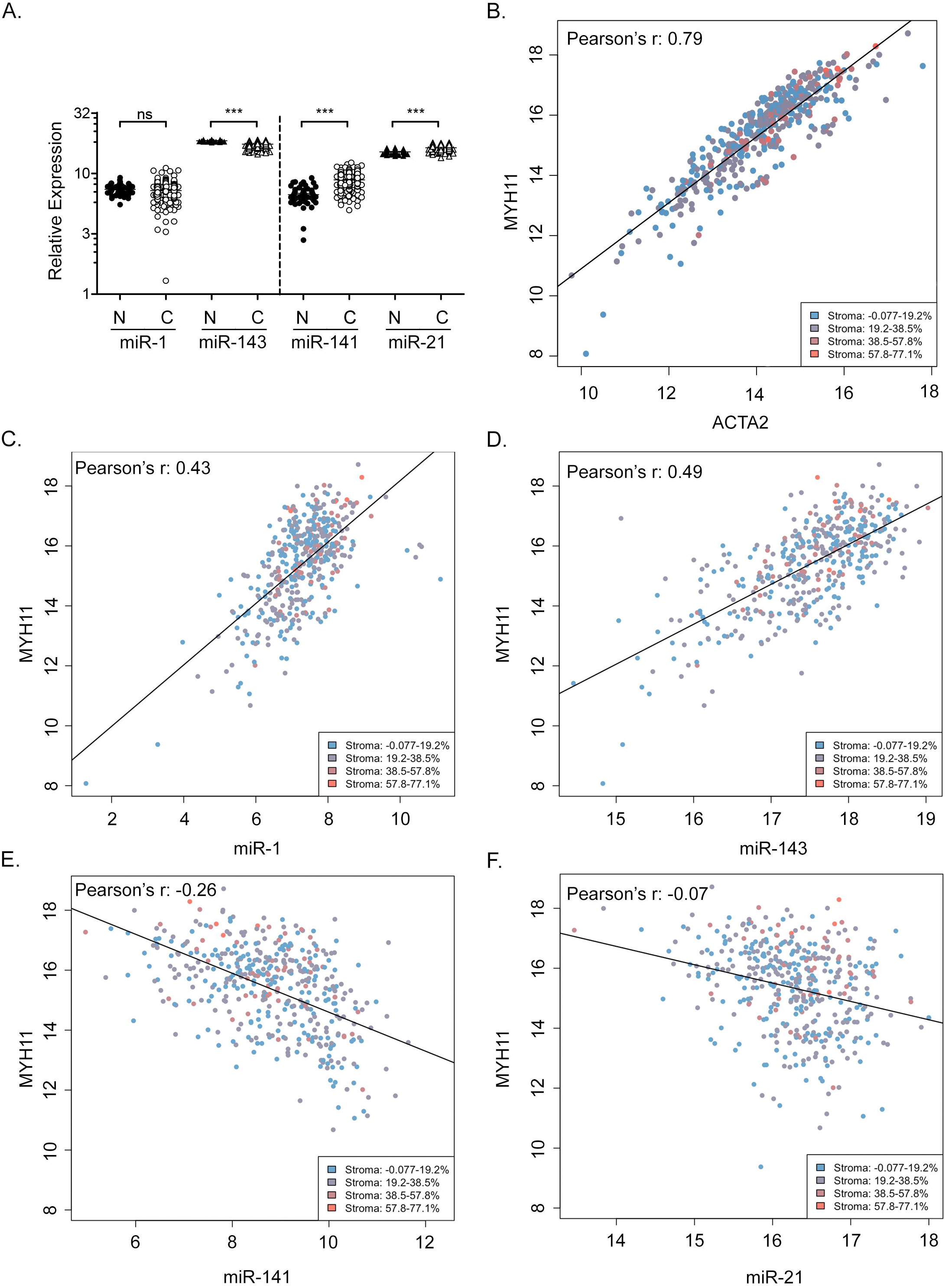
Cell-type selective miRNA expression in prostate cancer tissue from the TCGA. **A**. Relative miRNA expression levels in normal (N) prostate and PCa (C) tissue from TCGA-PRAD. Results are presented as mean and standard error. Log scale. P-values determined by Wilcox Rank Sum analysis. ns, not significant. ***, p < 0.001. **B**. Pearson’s correlation analysis of stromal markers MYH11 and ACT2 in TCGA-PRAD samples. p < 2.2 × 10^-16^. Legend indicates reported stromal content quadrants of adjacent sections. **C**. Pearson’s correlation analysis of miR-1 and stromal MYH11. p < 2.2 × 10^-16^. **D**. Pearson’s correlation analysis of miR-143 and stromal MYH11. p < 2.2 × 10^-16^. **E**. Pearson’s correlation analysis of miR-141 and stromal MYH11. p < 2.2 × 10^-16^. **F**. Pearson’s correlation analysis of miR-21 and stromal MYH11. p = 8.2 x 10^-9^. Pearson’s correlation coefficient determined by linear regression analysis and fitted using least-squares method.

To determine the cell-type specific nature of miRNA expression in this sample set, we considered correlations between each miRNA and two stromal mRNA markers, smooth muscle myosin heavy chain 11 (MYH11) and α-SMA (ACTA2). Both stromal gene expression markers were highly correlated across this large sample set (Figure 6B), with Pearson’s r values of 0.79 (p < 2.2 □ 10^−16^). We also considered the reported stromal content of adjacent sections used for TCGA-PRAD gene expression analysis, which are indicated in Figure 6B-6F by color coding. The reported stromal content was only weakly associated with the levels of stromal marker expression (Peerason’s r = 0.032; Supplementary Figure S2). These results suggest that the amount of stroma quantified on the adjacent tissue sections does not reflect the true amount of stroma present in the specimens used for RNA preparation. We therefore focused on stromal markers ACTA2 and MYC11. We found a strong positive correlation between miR-1 expression and stromal MYH11 (Figure 6C), with a Pearson’s r value of 0.43 (p < 2.2 × 10^-16^). MYH11 also strongly and directly correlated with miR-143, with a Pearson’s r value of 0.49 (p < 2.2 × 10^-16^) (Figure 6D). Levels of miR-141 had a strong and inverse correlation with stromal MYH11, with a Pearson’s r value of −0.26 (p < 2.2 × 10^-16^), suggesting epithelial rather than stromal gene expression (Figure 6E). Finally, miR-21 demonstrated a weak inverse correlation with stromal MYH11 (Figure 6F), with a Pearson’s r value of −0.07 (p = 8.2 × 10^-9^). These data indicate a predominantly stromal expression pattern for miR-1 and miR-143, and a predominantly epithelial expression pattern for miR-141. Similar and significant correlations were observed for each miRNA when compared with stromal ACTA2 (data not shown).

### Reduced miR-1 and elevated miR-21 are associated with biochemical recurrence

With the new understanding of the cell-type specific expression of these miRNAs, we then analyzed their association with biochemical recurrence and clinical and pathologic variables within the TCGA-PRAD dataset. In univariate analysis, Gleason grade (primary pattern G2 and G3 vs. G4 and G5), tumor stage (T2 vs. T3 and T4), lymph-node status (N0 vs. N1), and the dichotomized expression (using optimal thresholds) of all analyzed microRNAs were significantly associated with biochemical recurrence (Table 1A). Only miR-1 and miR-21 showed an association with biochemical recurrence as continuous variables. However, these pathologic features may also reflect differences in epithelial or stromal cell density (31). Therefore, multivariate analyses were also performed. Strikingly, elevated miR-21 levels were significantly associated with biochemical recurrence on top and beyond Gleason grade and stage in multivariate analysis (Table 1B, p = 0.044). Reduced miR-1 was also closely related to disease recurrence in multivariate analyses, but without significance (Table 1B, p = 0.055). These results suggest that loss of miR-1 and gain of miR-21 expression are highly associated with PCa aggressiveness and Gleason Score.

**Table 1 -.** Univariate and Multivariate Cox proportional hazard model on TCGA-PRAD miRNA levels and biochemical recurrence.

| **Predictors** | **HR** | **HR CI: 2.5%** | **HR CI: 97.5%** | **P-value** |
| --- | --- | --- | --- | --- |
| **(A) Univariate Analysis** |  |  |  |  |
| Primary Gleason (G2-G3 vs. G4-G5) | 3.275 | 0.582 | 1.791 | 0.000 |
| Stage (T2 vs. T3-T4) | 5.943 | 0.853 | 2.712 | 0.000 |
| Lymphnode status (N0 vs. N1) | 2.014 | 0.07 | 1.33 | 0.029 |
| Stromal content (%) | 1.002 | -0.017 | 0.02 | 0.852 |
| miR-21 expression (continuous) | 2.153 | 0.282 | 1.252 | 0.002 |
| miR-21 expression (dichotomous) | 2.796 | 0.475 | 1.581 | 0.000 |
| miR-1 expression (continuous) | 0.699 | -0.636 | -0.08 | 0.012 |
| miR-1 expression (dichotomous) | 0.245 | -2.259 | -0.555 | 0.001 |
| miR-141 (continuous) | 1.018 | -0.216 | 0.251 | 0.883 |
| miR-141 expression (dichotomous) | 0.39 | -1.87 | -0.013 | 0.047 |
| miR-143 (continuous) | 0.93 | -0.402 | 0.256 | 0.666 |
| miR-143 expression (dichotomous) | 0.416 | -1.799 | 0.048 | 0.063 |
| **(B) Multivariate Analysis** |  |  |  |  |
| **Primary Gleason + Stage** |  |  |  |  |
| Primary Gleason (G2-G3 vs. G4-G5) | 2.227 | 0.158 | 1.443 | 0.015 |
| Stage (T2 vs. T3-T4) | 3.519 | 0.272 | 2.245 | 0.012 |
| **Primary Gleason + Lymphnode status** |  |  |  |  |
| Primary Gleason (G2-G3 vs. G4-G5) | 3.234 | 0.499 | 1.849 | 0.001 |
| Lymphnode status (N0 vs. N1) | 1.439 | -0.271 | 0.999 | 0.261 |
| **Primary Gleason + Stage + Lymphnode status** |  |  |  |  |
| Primary Gleason (G2-G3 vs. G4-G5) | 2.131 | 0.053 | 1.46 | 0.035 |
| Stage (T2 vs. T3-T4) | 4.136 | 0.317 | 2.522 | 0.012 |
| Lymphnode status (N0 vs. N1) | 1.272 | -0.399 | 0.881 | 0.461 |
| **Primary Gleason + Stage + Lymphnode status + miR-21** |  |  |  |  |
| Primary Gleason (G2-G3 vs. G4-G5) | 1.964 | -0.042 | 1.392 | 0.065 |
| Stage (T2 vs. T3-T4) | 3.976 | 0.264 | 2.497 | 0.015 |
| Lymphnode status (N0 vs. N1) | 1.192 | -0.471 | 0.822 | 0.595 |
| miR-21 expression (dichotomous) | 1.816 | 0.015 | 1.178 | 0.044 |
| **Primary Gleason + Stage + Lymphnode status + miR-1** |  |  |  |  |
| Primary Gleason (G2-G3 vs. G4-G5) | 1.843 | -0.099 | 1.322 | 0.092 |
| Stage (T2 vs. T3-T4) | 3.734 | 0.215 | 2.42 | 0.019 |
| Lymphnode status (N0 vs. N1) | 1.127 | -0.525 | 0.765 | 0.716 |
| miR-1 expression (dichotomous) | 0.392 | -1.896 | 0.021 | 0.055 |
| **Primary Gleason + Stage + miR-1 + miR-21** |  |  |  |  |
| Primary Gleason (G2-G3 vs. G4-G5) | 1.824 | -0.06 | 1.262 | 0.075 |
| Stage (T2 vs. T3-T4) | 2.978 | 0.09 | 2.093 | 0.033 |
| miR-1 expression (dichotomous) | 0.478 | -1.635 | 0.159 | 0.107 |
| miR-21 expression (dichotomous) | 1.754 | -0.015 | 1.139 | 0.056 |

## DISCUSSION

Early gene expression profiles exhibited a clear association between miRNA levels, tumor developmental lineage, and differentiation state (1). These signatures, from whole tumor lysates, are reflective of miRNA expression from cancer cells and cancer-associated benign epithelium, vasculature, stroma, and immune infiltrates. This cellular heterogeneity can contribute to inconsistent conclusions between different miRNA gene expression studies (32). Therefore, there is a need to improve our understanding of cell-type specific miRNA expression patterns, and how they may impact the interpretation of miRNA function in tumor biology.

This may be best exemplified by the miR-143/145 gene family (4). A recent study of miR-143/145 in murine intestinal regeneration found that tissue repair was only compromised when miR-143/145 was specifically deleted from the mesenchyme, but not the epithelium. *In situ* hybridization and microdissection analyses further demonstrated near-exclusive miR-143/145 expression in murine and human intestinal mesenchyme, with undetectable expression in colon tumor epithelium (2). In a separate study of murine lung tumorigenesis, Dimitrova and colleagues found no significant changes in disease phenotype when miR-143/145 was deleted from lung epithelium (3). On the other hand, depletion of miR-143/145 from the tumor stromal microenvironment resulted in reduced tumorigenesis and diminished neoangiogenesis. Collectively, these studies implicate a stromal, rather than epithelial, role for miR-143/145 in human disease.

Our current study of human prostate tissues substantiates a predominantly stromal gene expression pattern for miR-143. Through two independent sources of microdissected tissue, we found that miR-143 is predominantly expressed within the prostatic stroma. Analyses of the TCGA dataset further support a direct correlation between miR-143 expression and the expression of stromal markers MYH11 and ACTA2. This new data, and those observed in colon and lung cancer, indicate that the loss of miR-143 in macrodissected tumors may due to differential stromal sampling between benign and malignant tissues. Our direct comparisons between normal and cancer stroma provide new data that miR-143 can also be specifically diminished in tumor-associated stroma. Thus, miR-143 expression may be specifically suppressed in the tumor microenvironment, or different cell types with altered miR-143 expression may be recruited to the tumor. We should note that miR-143 expression may not be exclusively stromal in these tissues. Very low levels of miR-143 could be detected in epithelium, and from pure populations of epithelial and cancer cell lines. For example, in some LNCaP cell extracts 3.9 copies of miR-143 could be detected, while in others, miR-143 was undetectable. This indicates that miR-143 expression is likely very low in epithelial cells, and near the limit of detection for some assays. From these data we conclude that miR-143 is predominantly, but not exclusively, stromal. This is consistent with the detection of miR-143 in multiple tissue and cell types in the SRA database.

MiR-1 has also been termed a tumor suppressor in prostate and other cancers (33–36). However, our results indicate that miR-1 is not likely a cell-autonomous tumor suppressor for PCa. We found miR-1 expression to be predominantly stromal in microdissected tissues. This result was supported by a clear association of miR-1 with stromal gene markers in the TCGA, and with little to no expression detected in epithelial cell lines. Interestingly, miR-1 expression was found in tissues microdissected by xMD, and not by LCM. These results suggest that different cell types may have been captured by xMD, versus LCM, or that there were differences in RNA quality. Indeed, miR-1 expression was extremely low, or absent, in fibroblast and fat samples from the SRA. The most significant miR-1 levels appear to be in human muscle, which is consistent with early studies of miR-1 expression and function in mice (37,38). Like miR-143, the levels of miR-1 were selectively decreased in the tumor-associated stroma of microdissected tissues. Thus, we posit that miR-1 expression is predominantly stromal, and that its expression may be suppressed in the tumor microenvironment. Alternatively, different cell types with altered miR-1 expression may populate the tumor microenvirontment.

MiR-141 was initially reported as a potential diagnostic serum biomarker for PCa (15). Consistent with these results, we found miR-141 to be significantly elevated in PCa. In contrast to miR-1 and miR-143, miR-141 expression was predominantly epithelial in human tissues and cell cultures. Extremely low levels of miR-141 could be detected in stromal tissue and cell cultures, indicating that miR-141 expression is predominantly, but not exclusively, epithelial. Numerous studies have also found elevated miR-21 expression in human PCa (8,17,39). Here we found elevated miR-21 expression in all PCa sample sets studied. Interestingly, our results indicate that miR-21 expression is highest in prostate stroma, rather than epithelium. This was consistent in normal prostate, as well as PCa, tissue. Previous studies have found elevated miR-21 expression in PCa-associated stroma, where it was associated with biochemical recurrence (40). Elevated stromal miR-21 expression has been similarly observed in other cancers (41,42). Unfortunately, we were not able to detect significant stromal- or epithelial-specific changes in miR-21 or miR-141 in microdisected PCa. This may indicate that the expression of these miRNAs is altered in both tumor stroma and epithelium, or that our sample size was too small to detect these differences in microdissected tissue. Significant differences could only be found for miR-1 and miR-143 in tumor-associated stroma in this small sample set. Larger studies may clarify these cell-specific expression patterns.

MiRNAs may be useful as biomarkers for PCa (43). Each of the four miRNAs evaluated in this study have been previously found to be associated with PCa disease aggressiveness, recurrence, or patient survival (34,40,44,45); however, their value as individual biomarkers remains unclear. Here, we found the dichotomized expression (using optimal thresholds) of all four microRNAs to be significantly associated with biochemical recurrence in the TCGA-PRAD dataset. However, only miR-1 and miR-21 showed an association with biochemical recurrence as continuous variables, and only miR-21 showed an association on top of and beyond Gleason grade and stage in multivariate analysis. It is notable that we previously did not find an association between miR-21 and biochemical recurrence in a smaller sample set 118 cases and controls matched for age, race, pathologic stage, and Gleason grade (46). Thus, with the larger TCGA-PRAD dataset, it appears that miR-21 levels may provide additional prognostic value beyond some clinical parameters; however, the benefits are likely moderate and may not be sufficient to improve clinical practice. Interestingly, miR-1 levels were nearly associated with disease recurrence in multivariate analyses (p = 0.055), suggesting that stromal miRNA expression may be informative for PCa prognoses. Further studies are needed to make this conclusion.

In summary, we have demonstrated for the first time predominant stromal expression of miR-1 and miR-143 in prostate tissue, predominant epithelial expression of miR-141, and both stromal and epithelial expression of miR-21. The specific and significant loss of miR-1 and miR-143 in tumor-associated stroma indicates a potential role for these miRNAs in the tumor microenvironment. The revelation of compartment-specific miRNA expression also sheds new light on the potential for each of these miRNAs as therapeutic targets and as biomarkers for determining disease progression.

## ACKNOWLEDGEMENTS

We thank Charles Ewing and William Isaacs for providing PrEC cells, Robert Veltri for providing RNA from BPH1 cells, and John Isaacs for discussions regarding prostatic stroma. We would also like to acknowledge the Tissue Services Core supported by the SKCCC CCSG (P30 CA006973) for their services and assistance, in addition to acknowledging the use of tissues procured by the National Disease Research Interchange (NDRI) with support from NIH grant 2 U42 OD011158.

